# Toolkit for Oscillatory Real-time Tracking and Estimation (TORTE)

**DOI:** 10.1101/2021.06.21.449019

**Authors:** Mark J Schatza, Ethan B Blackwood, Sumedh S Nagrale, Alik S Widge

## Abstract

**Background:** Closing the loop between brain activity and behavior is one of the most active areas of development in neuroscience. There is particular interest in developing closed-loop control of neural oscillations. Many studies report correlations between oscillations and functional processes. Oscillation-informed closed-loop experiments might determine whether these relationships are causal and would provide important mechanistic insights which may lead to new therapeutic tools. These closed-loop perturbations require accurate estimates of oscillatory phase and amplitude, which are challenging to compute in real time.

**New Method:** We developed an easy to implement, fast and accurate Toolkit for Oscillatory Real-time Tracking and Estimation (TORTE). TORTE operates with the open-source Open Ephys GUI (OEGUI) system, making it immediately compatible with a wide range of acquisition systems and experimental preparations.

**Results:** TORTE efficiently extracts oscillatory phase and amplitude from a target signal and includes a variety of options to trigger closed-loop perturbations. Implementing these tools into existing experiments is easy and adds minimal latency to existing protocols.

**Comparison with Existing Methods:** Most labs use in-house lab-specific approaches, limiting replication and extension of their experiments by other groups. Accuracy of the extracted analytic signal and accuracy of oscillation-informed perturbations with TORTE match presented results by these groups. However, TORTE provides access to these tools in a flexible, easy to use toolkit without requiring proprietary software.

**Conclusion:** We hope that that the availability of a high-quality, open-source, and broadly applicable toolkit will increase the number of labs able to perform oscillatory closed-loop experiments, and will improve the replicability of protocols and data across labs.

**Highlights:**

- TORTE provides a toolkit to investigate closed loop oscillation-informed experiments.
- The toolkit is versatile and open-source promoting replicability across scientists.
- The analytic signal algorithm within TORTE preforms equally to existing algorithms.

## 1. Introduction

### 1.1. Importance of Oscillations

Oscillations in continuous neural data are implicated in a wide range of functional processes, including decision making, learning and memory, sensory coordination, and emotion regulation. Dominant theories argue that cross-regional oscillatory synchrony (phase-phase and/or phase-amplitude coupling) enables and may be necessary for inter-regional communication (Engel et al., 2001; Buzsáki et al., 1994). That causal model has not yet been proven, as most prior work only shows correlations between oscillations, synchrony, and behavior. There is still a strong possibility that oscillations have no causal role, but are solely epiphenomena of spike-level processes (Schneider et al., 2020; Wilson et al., 2018; Tort et al., 2018). On the other hand, some early results suggest that oscillation-informed perturbations can alter brain circuit function in ways that are not possible with oscillation-blind approaches. Phase-locked stimulation can induce plasticity (Zanos et al., 2018; Zrenner et al., 2018), as can stimulation optimized to interact with a dominant cross-regional oscillation (Lo et al., 2020). Stimulation locked to the amplitude of a tremor-related oscillation can be more efficient in suppressing that tremor (Rosin et al., 2011; Bronte-Stewart et al., 2009), as can stimulation delivered at specific phases of a tremor cycle (Cagnan et al., 2019). Similar phase-aware approaches may be useful in manipulating circuits relevant to psychiatric illness (Herman and Widge, 2019; Widge and Miller, 2019; Kanta et al., 2019; Knudsen and Wallis, 2020).

### 1.2. Difficulty in Creating Closed Loop Experiments

Early closed-loop results are promising but highlight a major challenge in broadly testing causal claims about oscillatory synchrony – the need for accurate real-time estimates of an oscillation’s state. To demonstrate that cross-region or within-region phase-phase, or phase-amplitude, phenomena are causally linked to a functional process, neuroscientists and clinicians need tools to perturb those phenomena and/or to lock stimuli to specific oscillatory events. The standard algorithm to extract oscillatory features, the Hilbert transform (Cohen, 2014), is not well suited to a real-time situation. A Hilbert transform requires a large window of data around the time of interest, or else edge effects appear in its output. Individual labs have addressed this problem by developing special-purpose hardware and/or alternate algorithms, each with limitations. Some approaches have estimation inaccuracies that are too large to provide consistent results (Siegle and Wilson, 2014). Others are accurate but require special hardware that is not easily maintained without dedicated engineering staff. They may be prohibited by high cost and may be difficult to implement into existing experimental protocols (Kanta et al., 2019; Rodriguez Rivero and Ditterich, 2021; Escobar Sanabria et al., 2020; Zrenner et al., 2018; Shirinpour et al., 2020). Many solutions are built atop proprietary, closed-source software such as MATLAB and its toolboxes (Hassan et al., 2020; Zelmann et al., 2020). These factors greatly limit reproducibility. Further, many existing systems are only capable of identifying peak or trough phases of an ongoing oscillation (Siegle and Wilson, 2014; Rodriguez Rivero and Ditterich, 2021). They cannot support other paradigms such as detecting intermediate phases (Zanos et al., 2018), estimating phase response curves (Ermentrout et al., 2012; Holt et al., 2014) or oscillatory amplitude. There is a need for a toolkit that provides accurate oscillatory calculations in a pure software solution (to maximize flexibility) and that can readily be implemented in many labs and experimental settings.

### 1.3. Introducing TORTE

Here we provide a Toolkit for Oscillatory Real-time Tracking and Estimation (TORTE) that enables closed-loop oscillatory experiments. This toolkit implements a real-time algorithm to extract the analytic signal of the continuous neural data and is built within the Open Ephys GUI (OEGUI) system (Siegle et al., 2017). We developed TORTE with the intention of providing an easy to use, flexible and accurate system for scientists across a broad range of disciplines. OEGUI is interoperable with a variety of recording setups commonly used for rodent and non-human primate experiments, supports next-generation high-density silicon probes such as Neuropixels, and has been integrated with both invasive and non-invasive human recordings (Schatza and Blackwood, 2020; Black et al., 2017). A single software processing chain usable across many different preparations could accelerate scientific progress, just as open-source neural analysis toolkits have improved both speed and reproducibility (Oostenveld et al., 2011; Boki et al., 2010).

TORTE enables closed-loop experiments where perturbations are locked to arbitrary values of either the oscillatory phase or amplitude of continuous neural data. Fig. 1A presents an example of locking an event to the 180° phase of the slow frequency component of a local field potential (LFP) recording. This event in slow wave oscillations during sleep was used to trigger an auditory stimulus which led to enhanced memory consolidation (Ngo et al., 2013). When paired with transcranial magnetic stimulation (TMS), this same event in the alpha band facilitated long term potentiation in humans (Zrenner et al., 2018). In Fig. 1B, we depict a perturbation being presented during a time when the oscillation is at a high amplitude. Using the amplitude of a motor cortical oscillation to trigger stimulation has been used to create a brain computer interface to restore motor function in nonhuman primates (Fetz, 2015), and in humans using the cortico-thalamic circuit to suppress tremor (Opri et al., 2020; Bronte-Stewart et al., 2009). These are exam-ples of two event targets and a few output stimuli. The versatility of TORTE allows users to target events for any phase or amplitude, at any frequency band in which true oscillatory activity occurs. This event can then be used to trigger any relevant perturbation, which may include presentation of a task stimulus, delivery of a reward/outcome, or direct brain electrical/optical/magnetic stimulation.

**Figure 1:**
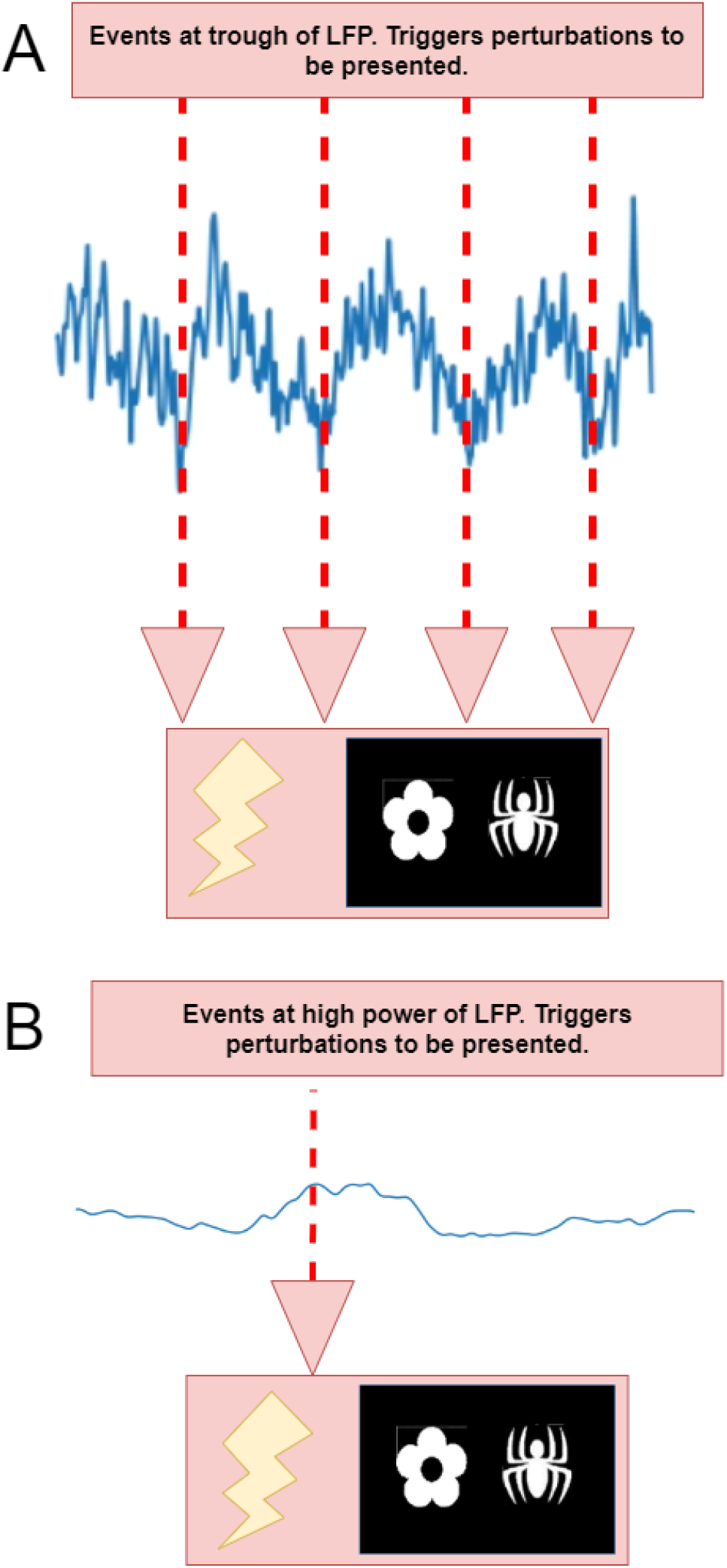
A) Events triggered at trough (180°) of low frequency oscillation. B) Event triggered by high power activity of a low frequency oscillation.

## 3. Materials and Methods

### 3.1. TORTE Overview

TORTE provides closed-loop tools to lock perturbations to neural oscillatory events. The toolkit, developed within OEGUI, can be run on any standard lab grade computer, including laptops for portable data acquisition, and can communicate with a variety of neural acquisition systems. The toolkit and OEGUI are freely available and modifiable. The GitHub repository includes extensive documentation on configuration and typical use cases (Schatza, 2021b).

Fig. 2 provides a system overview of the toolkit, highlighting the three main components. The “Open Ephys System” extracts oscillatory features from the neural data (Fig. 2A). The “Closed-Loop Hardware” layer handles communication between the experiment and OEGUI (Fig. 2B). The “Experiment” includes the subject and the presented/delivered perturbations (Fig. 2C). Starting with closed-loop hardware, an acquisition system records continuous neural data. This data is brought into OEGUI by a data interface plugin. Plugins currently available include: Open Ephys acquisition box, Alpha Omega intraoperative monitoring systems, Neuralynx systems, Neuropixels, EEG via a custom interface board (Black et al., 2017), and EEG using the LSL-inlet plugin (Schatza, 2021a). Additional systems are occasionally being added. Collectively, these systems cover common platforms for human, non-human primate, and rodent recordings. The neural data is passed on to the Real-Time Analytic Signal plugin. This plugin outputs either the phase or amplitude of the signal. The Analytic Signal Crossing Detector plugin continuously monitors the output from the Real-Time Analytic Signal plugin and triggers events when a threshold is crossed. The threshold is set to either a specific phase of interest or an amplitude value. Additional logic can denoise these signals if needed, e.g., by requiring the amplitude to cross and remain on one side of a threshold for M of N samples. A rudimentary artifact suppression algorithm is included that limits the jump size between samples. We have not, at the present time, implemented more advanced approaches such as stimulation artifact template subtraction. These would likely best be achieved by separate plugins, to maximize use of OEGUI’s modular design. TORTE is best used for situations where the presented perturbation does not induce artifact (e.g., optical or sensory stimulation with electrical recordings), or in situations where a long recovery time can be given between pulses for amplifiers to settle. The Event Broadcaster plugin then uses the common, very low latency interprocess communication framework ZeroMQ (0MQ, 2021) to output the Crossing Detector event to the closed-loop hardware using a publisher/subscriber mode of communication. Any of the 26 programming languages that ZeroMQ supports can be used to create output logic to receive this event. Output logic code is not provided within this toolkit, as it is heavily dependent on the specific perturbation to be delivered. However, example code that receives ZeroMQ events in Python can be found in the provided OEGUI Python tools repository (Siegle, 2017) and LabVIEW code can be found on our GitHub (Blackwood, 2021). We specifically include an example of how to generate a 5 V rising-edge square pulse, the most common signal used to trigger both brain stimulation and task-related hardware, but the output logic can be designed to create stimuli of any kind. Further, ZeroMQ supports communication within a single computer (e.g., for controlling physiology and direct brain stimulation in closedloop) or network communication (e.g., for synchronizing multiple experimental machines via standard Internet protocols). This decoupling of detection from output allows easier incorporation of this toolkit into existing experimental protocols. For versatility and computational efficiency OEGUI and TORTE are C++ based. To access the toolkit the code can either be compiled from source for advanced users, or simply installed using binaries provided across all major platforms.

**Figure 2:**
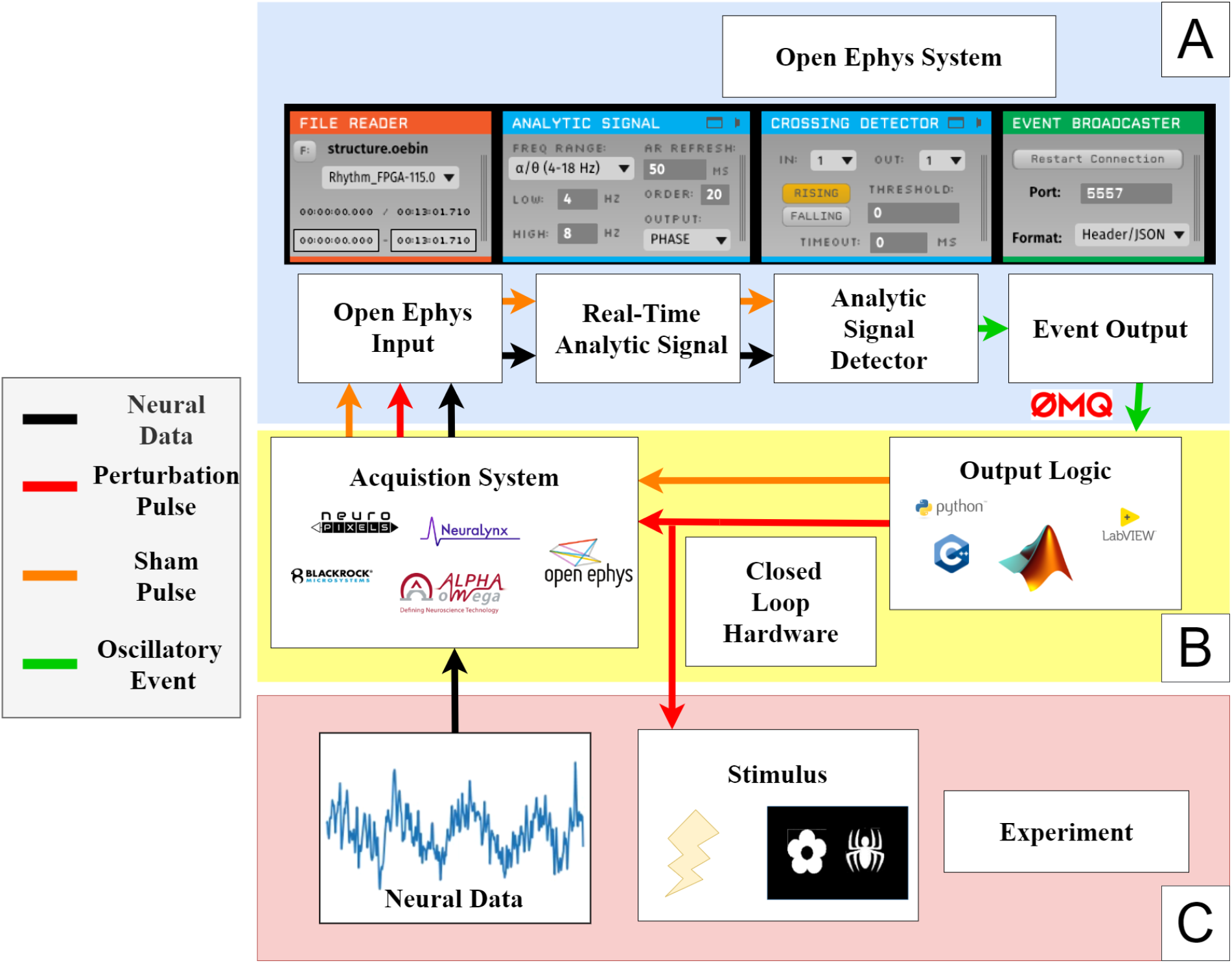
A) Overview of the software (Open Ephys System). B) Closed-loop hardware of the system includes an acquisition system and output logic (Closed-Loop Hardware). C) The experiment includes the subject and the experimental stimuli presented to them (Experiment).

To improve accuracy in phase-locked closed-loop experiments and improve data analysis, it is recommended to provide feedback of perturbation timing back into the system. There are a few methods to implement this feedback. It is recommended to provide either a digital input into the system or send a software TTL pulse to OEGUI with ZeroMQ. If neither of these are possible, an alternative technique would be to send perturbation event markers through an analog input (e.g., by consuming one recording channel). Two types of pulses are typically sent, a perturbation pulse (Fig. 2 red arrow) and a sham pulse (Fig. 2 orange arrow). The perturbation pulse is tightly time-locked to the simultaneously acquired neural data and can be used in later analysis to extract data at the time of perturbation. The sham pulse can be used to improve accuracy in phase-locked closed-loop experiments in real time and/or used to verify the oscillations’ status during the event trigger timing without perturbation related artifacts. The sham pulse is sent in place of presenting a perturbation, but with the same timing of a perturbation pulse. If used to improve accuracy in phase-locked closed-loop in real time, the Analytic Signal Plugin and Analytic Signal Detector are set up to listen for these events. As described further below, this enables a self-adjusting algorithm that compensates for experimental hardware latency and bias in phase estimates.

### 2.2. Analytic Signal Calculation

TORTE transforms continuous neural data into its analytic signal in real-time utilizing a Hilbert transformer. The Analytic Signal Plugin GUI allows a user to adjust the algorithm for their experiment. Fig. 3 shows the flowchart for the Hilbert transformer (Fig. 3A), how the customizable values on the plugin’s user interface affects the algorithm (Fig. 3B), and the corresponding output (Fig. 3C).

**Figure 3:**
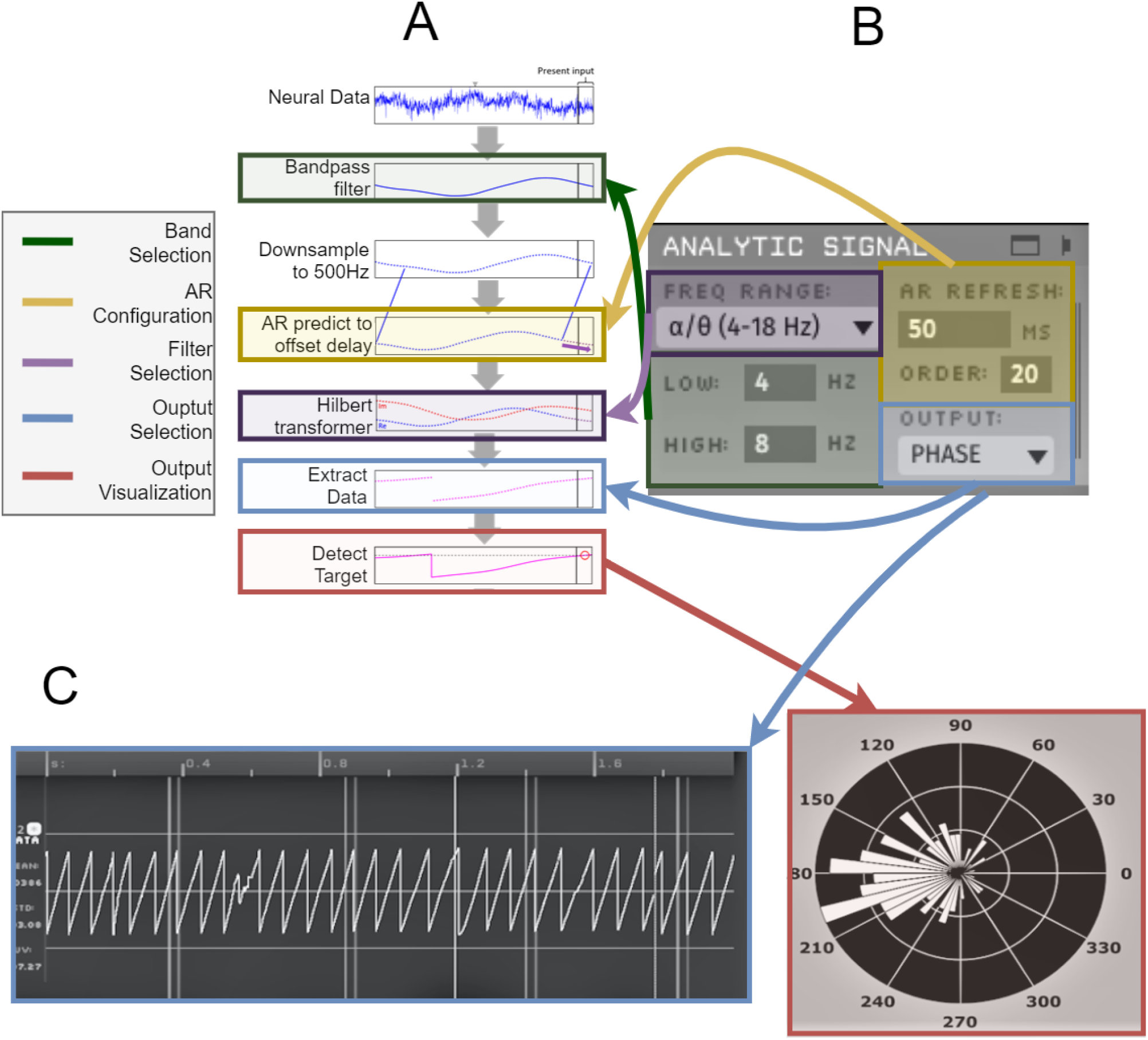
A) Flow chart overviewing the algorithm transforming neural data into oscillatory activity. B) GUI for the analytic signal plugin. C) GUI elements showing output of analytic signal plugin.

Fig. 3A provides a flow chart of the algorithm. It starts with raw continuous neural data, e.g., a single EEG or LFP channel. This may be a derived channel, e.g., a bipolar-referenced pair as in (Zelmann et al., 2020) or local Laplacian as in (Zrenner et al., 2018). This data is causally filtered through a 2nd order forward Butterworth bandpass filter to extract a frequency of interest. For the next steps of the algorithm, the data is downsampled to 500 Hz. An autoregressive (AR) model then predicts enough samples ahead in time to compensate for the Hilbert transformer’s group delay, as described below, similar in principle to (Blackwood et al., 2018). The AR order and model refresh rate for computing the AR model coefficients can be configured by the user to achieve the optimal efficiency/accuracy tradeoff for their system. With default parameters, the model coefficients are computed using the last 1 second of data and are updated every 50 ms for a 20th order AR model. Decreasing the update frequency and the order of the model can greatly improve computational efficiency. It is expected that no artifacts are present within the data used for the AR prediction during event detection. With default parameters this would be 20 samples at 500 Hz or 40 ms of data. To ensure no artifacts are present, the user should enable a timeout/lockout period between successive detections and perturbations. This is an included feature of the TORTE plugins. A Hilbert transformer is then applied to the predicted and observed values, returning the imaginary component at quadrature with the data. The Hilbert transformer is a finite impulse response (FIR) filter that has a phase response with a constant group delay (offset) equal to half the filter order. The AR model predicts the bandpassed signal sufficiently far into the future to compensate for this delay. TORTE currently includes five Hilbert transformers that provide well-behaved amplitude responses that are close to flat in the band of interest and are reasonably flat and suppressed outside the band of interest. Filters are provided for oscillatory bands of alpha/theta (4-18 Hz), beta (10-40 Hz), low gamma (30-55 Hz), mid gamma (40-90 Hz), and high gamma (60-120 Hz). Users can add new filters that provide a better response for their band of interest into the plugin. This could be used to extend TORTE for detection of, e.g., sharp wave ripples near 200 Hz. Instructions are provided in the TORTE GitHub repository to implement a new filter, which is trivial to add to the plugin once the filter has been created. Parks-McClellan optimal FIR filter design, e.g., MATLAB’s firpm or Python’s scipy.signal.remez function, is generally adequate. The algorithm downsamples to 500 Hz so filters up to 250 Hz can easily be created. The downsampling frequency was chosen as it provides good tradeoffs for efficiency, accuracy and range of frequencies. Motivated users could adjust this sampling rate for highly specific needs, but it is not an easy to adjust feature of this toolkit.

The band selection region (green) of the phase calculator can be used for updating the frequency band of interest for the initial bandpass. The filter selection region (purple) is used to choose which of the Hilbert transformers best fits the frequency band of interest. The AR configuration region (yellow) is used to adjust the efficiency/accuracy tradeoff for the AR model. The output selection region (blue) allows the user to select either phase or amplitude for the output. The output visualization region (red) shows real time accuracy if using phase-locked closed-loop.

TORTE was compared to the standard peak/trough algorithm built into OEGUI (Siegle et al., 2017). The standard algorithm can compute phase, but notably cannot compute amplitude, of a target signal. Further, it can only detect 0, 90, 180, or 270° phase events. To find the target, the standard algorithm bandpasses the data down to the frequency of interest and then detects zero crossings or slope inversions. Further optimization of this type of algorithm is possible, but the standard algorithm tested in this publication is a direct representation of the next best option in the current Open Ephys environment.

### 2.3. Learning Algorithm

TORTE is interoperable with a wide range of systems and output hardware. Each laboratory setup will have unique sources of latency between the triggering oscillatory event and the delivery of the matched perturbation. To improve phase-locked closed-loop experimentation and reduce the effects of such delay, TORTE includes a learning algorithm. Fig. 4A shows an example intending to lock a perturbation to a phase of 180º, but with an expected communication latency that will produce an approximately 20° offset at the center frequency of the desired band. Thus, initially events are commanded to trigger when the phase crosses 160° to account for the offset. The learning algorithm will iteratively improve this offset throughout the experiment, to optimize perturbation delivery at the target phase. For common cases of electrical/optical stimulation that cause recording artifacts, learning requires sham pulses. For perturbations that do not cause artifacts (e.g., oscillation-locked delivery of sensory stimuli), the perturbation pulses may be used instead. Fig. 4B then shows learning, as implemented in the Analytic Signal Crossing Detector GUI. In this window, the event channel, target phase, learning rate and other parameters are configured. The event channel is set to track whatever source is receiving sham/perturbation pulse events. After the event is received, TORTE waits for additional neural data. It then performs an acausal calculation of phase using a bidirectional filter and full (not approximated) Hilbert transform. This acausal phase will be more accurate than the real time estimate. Using the acausal phase calculation, TORTE compares the phase at which the sham pulse arrived to the target phase. The circular difference between the phases is multiplied by the current learning rate to adjust the threshold value. The learning rate decays over time as configured by the user, and the process usually can converge in a few minutes.

**Figure 4:**
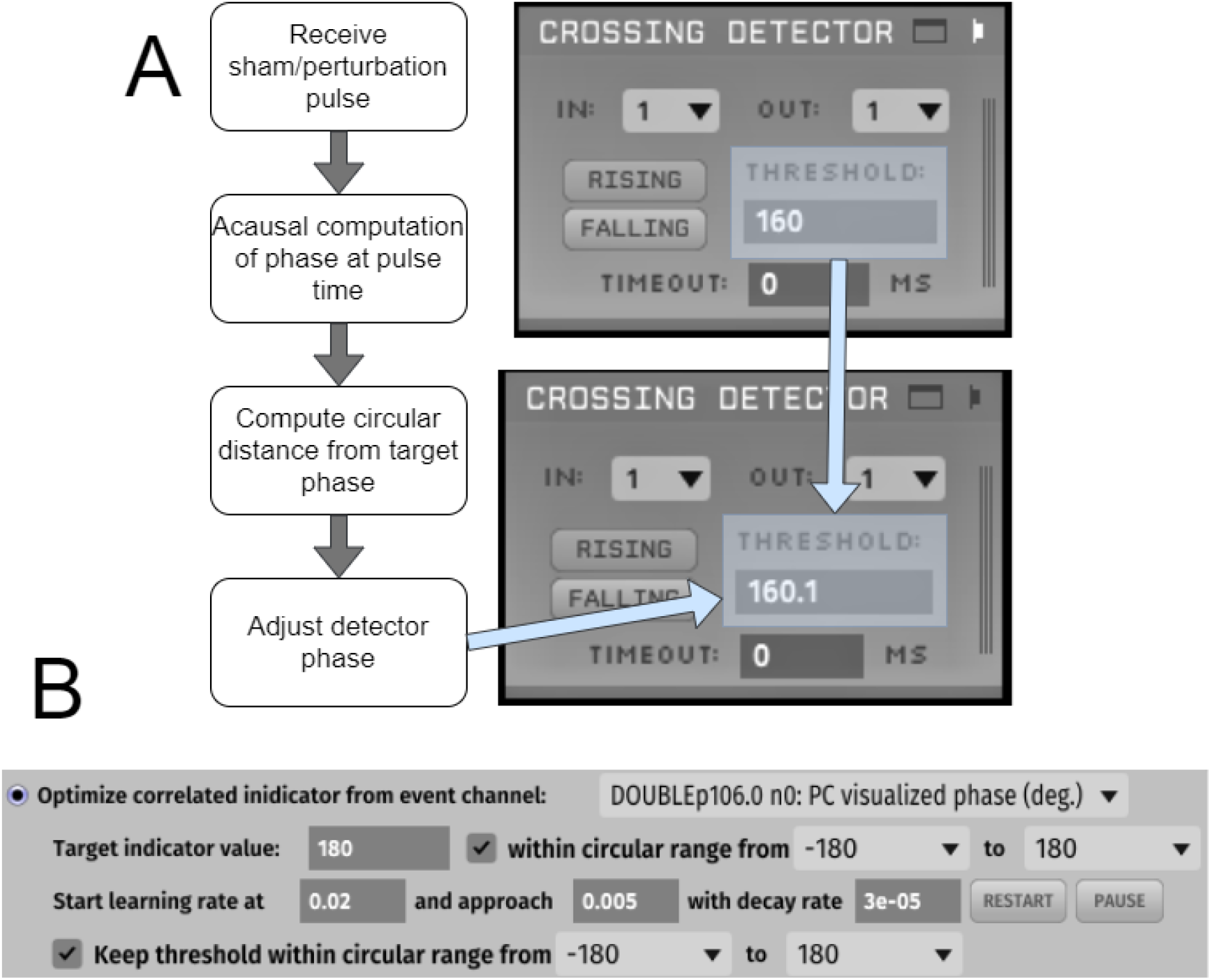
A) Flowchart of learning algorithm. Right side shows the crossing detector plugin being updated. Left side shows the logical flowchart for the learning algorithm. B) Crossing detector GUI showing variables conigurable for the learning algorithm.

### 2.4. Real-time Feedback of Coherence and Spectrogram

A likely use case for TORTE is closed-loop control of oscillations, and an experimenter may wish to verify that this control is effective as the experiment progresses. TORTE thus includes a Coherence and Spectrogram Viewer, which displays either the coherence between multiple channels or the spectrogram of individual channels. Both of these values can be determined from a time frequency representation (TFR) of the data. See Fig. 5A for a flowchart describing the TFR decomposition. The decomposition starts by storing data into a buffer of a size configured by the user. The default buffer size is 8 seconds, which provides a reasonable balance between estimation accuracy (number of oscillatory cycles contained in a buffer) and frequency of updates. Any buffer size above 4 seconds will provide reasonable calculations for both coherence and spectrogram at most frequency bands. Once the buffer is filled, a 2 second Hann window is used to perform TF decomposition using a sliding window Fourier transform. With the TFR calculated, the power and covariance (cross spectral) matrices for the buffer can be calculated. Because TFR windows are calculated in real time, the user can choose to either weight these matrices linearly over the entire experiment or can use exponential weighting to emphasize recent changes in activity. Channels, frequencies of interest, and weighting factors are all configurable in this plugin. With the TFR computed, the plugins can show either the coherence between two groups of channels, or the spectrogram of the channels selected.

**Figure 5:**
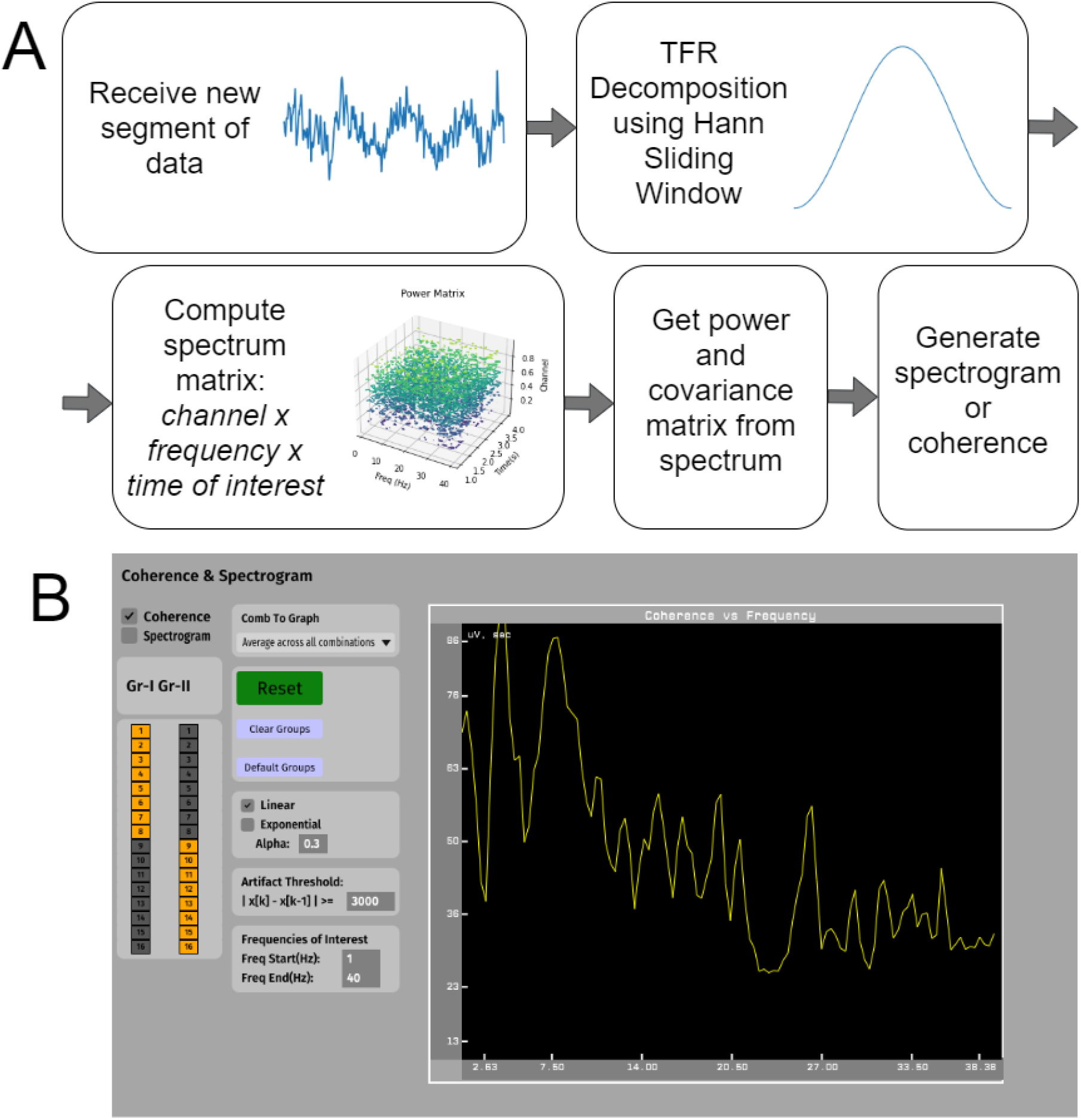
A) Flowchart describing calculation of real-time feedback using a TFR decomposition to generate either a spectrogram or coherence plot. B) Example screenshot of the UI for the coherence spectrogram visualizationl. The snapshot shown here is to demonstrate what a user may expect to see when using the plugin.

### 2.5. Experimental Validation

Four data sets were used to assess the algorithms described above. These datasets will be called Rodent, EEG, Simulated and Human. The Rodent dataset consists of 5 minute LFP recordings collected from the infralimbic cortex (IL) and basolateral amygdala (BLA) in freely behaving Long Evans rats (Lo et al., 2020). LFP were acquired continuously at 30 kHz (OpenEphys, Cambridge, MA, USA). An adaptor connected the recording head stage (RHD 2132, Intan Technologies LLC, Los Angeles, CA, USA) to two Millmax male-male connectors (8 channels each, Part number: ED90267-ND, Digi-Key Electronics, Thief River Falls, MN, USA). Sham and stimulation pulses were triggered at 180° for a 4-8 Hz oscillation. Pulses were commanded by a Python application which received events from an Open Ephys Event Broadcaster plugin. Sham pulses were used to assess accuracy post hoc. This dataset was procured in compliance with relevant laws and institutional guidelines; all animal procedures were reviewed and approved by the University of Minnesota IACUC. The EEG dataset consisted of two single channel 30 minute recordings located over the left prefrontal cortex (Brain Vision, Morrisville, NC, USA). This dataset was procured in compliance with relevant laws and institutional guidelines; all EEG recordings were reviewed and approved by the University of Minnesota IRB.

The Human dataset was collected from two individuals with refractory epilepsy (1 male) undergoing invasive monitoring as part of their clinical care at the Northwestern Memorial Hospital Comprehensive Epilepsy Center. sEEG depth electrodes (∼1 mm diameter, ∼2 mm contact length; AD-Tech Medical Instruments Co., Oak Creek, WI) were implanted according to clinical need prior to participation. Recordings were acquired using a Neuralynx ATLAS system with a scalp electrode reference and ground (Neuralynx, Boseman, MT). FIR digital bandpass filters were applied from 0.1 to 5000 Hz at the time of recording. Data were recorded at a resolution of 0.15 μV (5000 μV input range) and a sampling rate of 20 kHz, and were subsequently downsampled to 1 kHz for analysis. Data were streamed from the ATLAS to a separate computer running OEGUI via fiber optic cable. Sham and stimulation pulses were triggered at 180° for a 4-8 Hz oscillation through the OEGUI machine’s parallel port, using a custom OE plugin. Sham pulses were used to assess accuracy post hoc. The Simulated dataset was created in MATLAB by creating a sin wave with the frequency of interest with an amplitude that varies slightly over time. Pink noise was added on top of the sin wave with an amplitude equal to the peak amplitude from the frequency of interest. An example of a single Rodent and EEG dataset are available in the GitHub repository with Human data accessible by request.

For saline testing one channel of data within the infralimbic cortex of the Rodent dataset from each recording was converted into an MP3 file and played through an auxiliary port of the test computer with Audacity®. The exposed male end of the auxiliary cable was placed in a saline bath to emulate a brain. Recordings from the saline were then taken using the same headstage and electrodes as the in-vivo setup, but now measuring the replayed LFP data. In this testing setup, we did not deliver electrical stimuli or other perturbations back into the saline, but sent a 1 V rising-edge square pulse to the Open Ephys Acquisition system to track the timing of the detection events. To further characterize TORTE’s performance across a wide range of frequencies and conditions, and to collect more data points per condition, we further implemented a software simulation of this saline test. For the simulation, Rodent, EEG and Simulated datasets were loaded into MATLAB and processed by a MATLAB implementation of TORTE’s real-time buffered processing. System latencies between components were simulated by generating random numbers from a gamma distribution whose peak matched the median latency observed in the saline tests. We verified that the simulated saline test produced identical results to its physical counterpart. However, because data buffers could be “acquired” faster than real time in the simulation, we could complete each test thousands of times more quickly and use more datasets.

The MATLAB simulation was used to compare the TORTE and standard algorithm parameters by setting them to trigger events targeted at 180° and at 300° for oscillations ranging from 5 Hz to 55 Hz using the Simulated dataset. We chose 180° because it has been a target in practical closed-loop experiments (Zrenner et al., 2018) and 300° because it is not easily detectable by peak-trough or zero-crossing detectors. Phase-locked closed-loop experiments typically target lower frequency oscillations because they are commonly implicated in functional processes (Watrous et al., 2015) and their estimation is less affected by inherent system latencies. Using the MATLAB replication software, a phase-locked protocol was run targeting oscillations between 5 Hz and 55 Hz, with a step size of 1 Hz, for the two phase targets. The difference between the ground truth phase at the sham pulse time and the target phase was calculated. Ground truth phase was calculated using the standard offline approach of a forward-backward bandpass filter over the frequency band of interest, followed by a Hilbert transform. The TORTE Hilbert transformer method utilized its learning algorithm to iteratively improve the threshold for event triggers. The standard algorithm was set in trough (180°) mode for both targets; this highlighted certain features of that algorithm more clearly than setting it to 270° for the 300° target.

The MATLAB implementation was also used to simulate an in-vivo experiment in both the Rodent and EEG datasets to assess if any differences were present across recording techniques/species in both algorithms. The standard algorithm’s best parameters, targeting either 0, 90, 180 or 270°, were used on both algorithms and tests were completed targeting the theta band (4-8 Hz). Once again the learning algorithm was implemented for TORTE. To assess the impact on peak frequency within the oscillation of interest the spectrogram of the data across 1 minute windows was calculated using the pwelch function in MATLAB. The corresponding output was smoothed using a 5 minute gaussian window.

As a demonstration of how the overall system architecture and performance can vary depending on the specific acquisition hardware, and to further demonstrate TORTE’s viability for human closed-loop experiments, we performed a further test using an ATLAS human-grade electrophysiologic rig (Neuralynx, Bozeman, MT, USA). The Rodent data was played into the analog input of the ATLAS system using a USB DAQ and LabVIEW. The ATLAS system then broadcast the data as UDP packets over an Ethernet cable. A freely available acquisition plugin (Schatza and Blackwood, 2020) reassembled these packets as a datastream within OEGUI. On phase detection event triggers, a 1 V rising-edge square pulse was sent to the ATLAS system. ATLAS rebroadcasted the received square pulses alongside the data within the UDP packets.

All results were compared to a standard offline analysis procedure in MATLAB using Fieldtrip (Oostenveld et al., 2011). The MATLAB library CircStat (Berens, 2009) was used to determine circular mean and circular standard deviation (SD).

To assess the real-time feedback visualizations of power and coherence, the Rodent dataset was replayed in OEGUI using the file reader plugin with the IL and BLA channels split into separate groups for coherence measurements. The output for each trial was recorded by TORTE. The builtin functions included in Fieldtrip, ft_freqanalysis and ft_connectivityanalysis, were used to generate coherence and spectrogram outputs for the data at the same timepoints as the TORTE output from the Rodent data.

## 3. Results

### 3.1. Results Overview

In this section the efficacy of TORTE in a standard lab setup is shown. First we show that the system accurately estimates phase and amplitude. We then describe sources of latency within the system and how the online learning algorithm reduces latency effects. Finally, the coherence and power calculated by the casual monitoring algorithm are compared to an acausal MATLAB implementation to show adequate accuracy.

### 3.2. Phase Accuracy

Using the software replication experiment with the Simulated dataset, results for event phase accuracy at two phase targets for TORTE and the standard algorithm are shown in Fig. 6A-B and compiled in Table 1. TORTE triggers events (n=200000) within 2.4° of the target phase and with less than 41° SD at all frequencies and both phase targets. Targeting the trough, the standard algorithm triggers events with a mean error of 50° and a SD of 3° from the target phase at lower frequencies, with worsening performance as frequencies increase. While targeting 300° at the lowest frequency, the standard algorithm starts 70° from target with a SD of 3°. As the frequencies increase, the inaccuracy of the standard algorithm caused by the latency of the system makes the algorithm “accidentally” hit the target phase at around 40 Hz. The user could use this to their advantage, but would greatly limit target phase and frequency combinations.

**Table 1.** TORTE and Standard algorithm results from the Rodent dataset in saline and Simulated dataset in MATLAB.

|  | Saline<br>4-8 Hz 180° | MATLAB<br>5 Hz 180° | MATLAB<br>55 Hz 180° | MATLAB<br>5 Hz 300° | MATLAB<br>55 Hz 300° |
| --- | --- | --- | --- | --- | --- |
| Mean TORTE | 0.42° | -0.86° | -2.4° | -0.39° | -2.26° |
| SD TORTE | 63.5° | 16.42° | 40.53° | 11.9° | 40.98° |
| Mean Standard | -49.00° | -50.71° | -178.03° | 69.38° | -57.94° |
| SD Standard | 67.81° | 3.07° | 26.55° | 3.07° | 26.55° |

**Figure 6:**
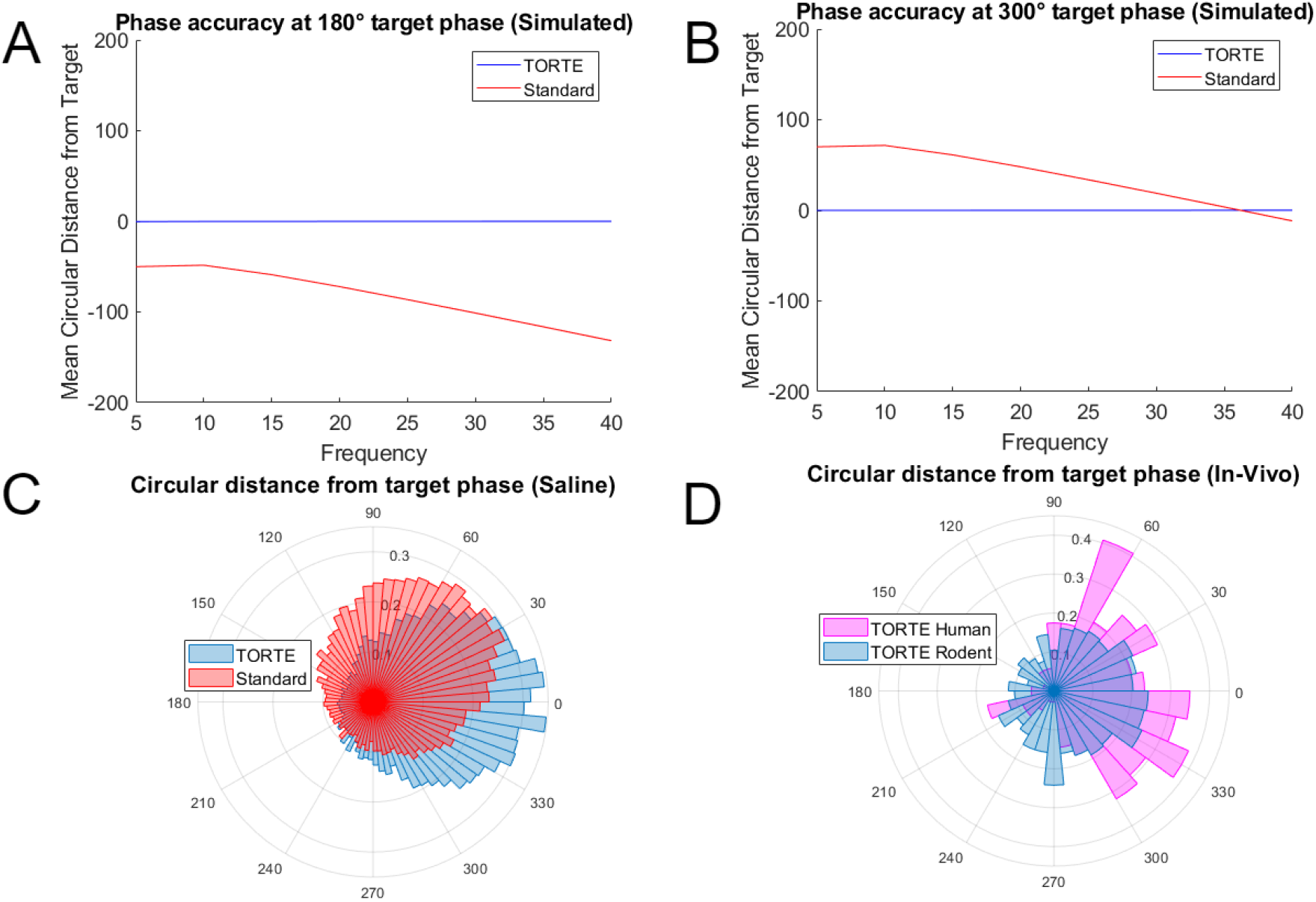
A) Comparison of circular distance from target phase in MATLAB across frequencies for TORTE and the standard algorithm targeting 180 and B) 300. Standard error of the mean is shown as a shaded region, but is too small to be seen in the figure. C) Circular distance from 180 phase target in the saline bath setup. D) Circular distance from target phase in Rodent and Human in-vivo experiments.

The saline bath phase-locked closed-loop experiments were set to target 180° phase within the 4-8 Hz band within the selected IL data channel (n=115). As seen in Fig. 6C a similar offset appears between the mean of the standard algorithm event phases and the target phase. In-vivo experiments with the same target are shown in Fig. 6D for both Rodent (n=2) and Human (n=2) closed loop experiments. Compiled results for the in-vivo experiments are found in Table 2 and are comparable to saline and software tests.

**Table 2.** TORTE results across recording types.

|  | TORTE Rodent (In-Vivo) | TORTE Human | TORTE EEG (MATLAB) |
| --- | --- | --- | --- |
| Mean | 0.42° | 2.4° | 0.50° |
| SD | 63.5° | 60.5° | 80.56° |

Further MATLAB software implementation experiments tested the standard algorithm’s best case targets of 0, 90, 180 and 270° for an oscillation of 4-8 Hz. These tests were performed on the Rodent (Fig. 7A) and EEG (Fig. 7C) datasets. The highest power frequency within the Rodent dataset was found for each minute of the recordings (Fig. 7B). The peak oscillation frequency varies during the experiment, but phase accuracy remains high. These three experiments all verify the Hilbert transformer algorithm as being more versatile and more accurate than the standard algorithm.

**Figure 7:**
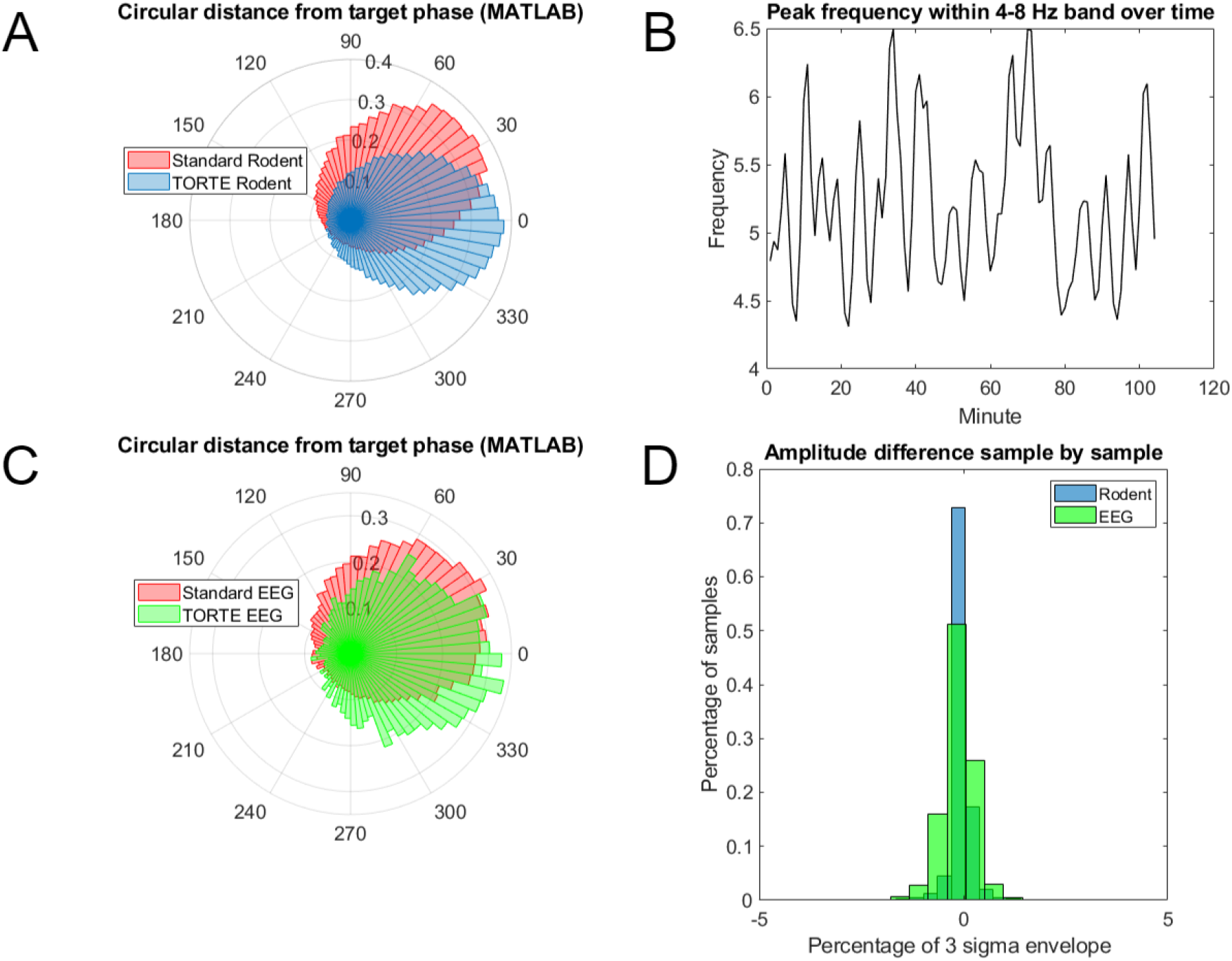
A) Comparison of circular distance from target phase in the theta band (4-8 Hz) using the MATLAB simulation for TORTE and the standard algorithm, targeting 0, 90, 180, and 270 in the Rodent dataset. B) Peak frequency within the theta band over time. C) Comparison of circular distance from target phase in the theta band using the MATLAB simulation for TORTE and the standard a gorithm, targeting 0, 90, 180, and 270° in the EEG dataset. E) ifference in real-time estimated amplitude to ground truth.

The TORTE results presented in Table 2 are also comparable to other approaches to high-accuracy phase prediction. (Zanos et al., 2018) reported an equivalent mean error to TORTE, but with a 30° higher standard deviation of phase accuracy. A similar AR prediction algorithm reported a mean error of 1° and a SD of 53° from their target phase (Zrenner et al., 2018). An alternative technique called Educated Temporal Prediction (ETP) has been proposed that estimates the phase oscillations and makes an educated guess at phase timings in the future. This method again yielded similar results, with a mean phase error of 0.37° and a SD of 67.35° (Shirinpour et al., 2020). Compiled results are shown in Table 3, emphasizing that the differences between each algorithm’s phase-locking performance are numerically quite small.

**Table 3.** Comparing reported results of state-of-the-art real-time phase estimates.

|  | TORTE | Zanos et al. (2018) | Zrenner et al. (2018) | Shirinpour et al. (2020) |
| --- | --- | --- | --- | --- |
| Mean | 0.42° | 41° | 1° | 0.37° |
| SD | 63.5° | 66° | 53° | 67.35° |

### 3.3. Amplitude Accuracy

Using the software replication setup with both the Rodent and EEG datasets, the real-time amplitude output from the TORTE Hilbert Transformer algorithm was recorded. The resulting output was compared sample by sample with an amplitude ground truth calculation using the standard offline Hilbert transform approach. Fig. 7D shows the difference in amplitude between both outputs as the percentage of the three sigma envelope of all amplitude values. Ampli-tude differences are minimal across the experiment for both datasets.

### 3.4. Latency

A wide variety of acquisition systems can provide data to the OEGUI, each with unique latency and jitter, ranging from μs to ms. As an example, we show how the latency of the Open Ephys acquisition board is driven by its USB-based communication, and compare this to the Ethernet-based Neuralynx ATLAS. Fig. 8A shows the latency between an event occurring in the neural data and a 1 V rising-edge square pulse being sent back to the preparation in response. The Ethernet-based system has a lower mean latency and much narrower spread. Fig. 8B shows the processing time of the TORTE algorithm for a buffer with 18.3ms of neural data. Using these two data sets, we can determine what percentage of the real-time closed-loop latency is attributable to the TORTE algorithms. TORTE’s internal calculations comprise about 0.9% of the latency in the Open Ephys acquisition system and 4% of the latency in the Neuralynx ATLAS, demonstrating that the majority of latency lies in inter-system communication.

**Figure 8:**
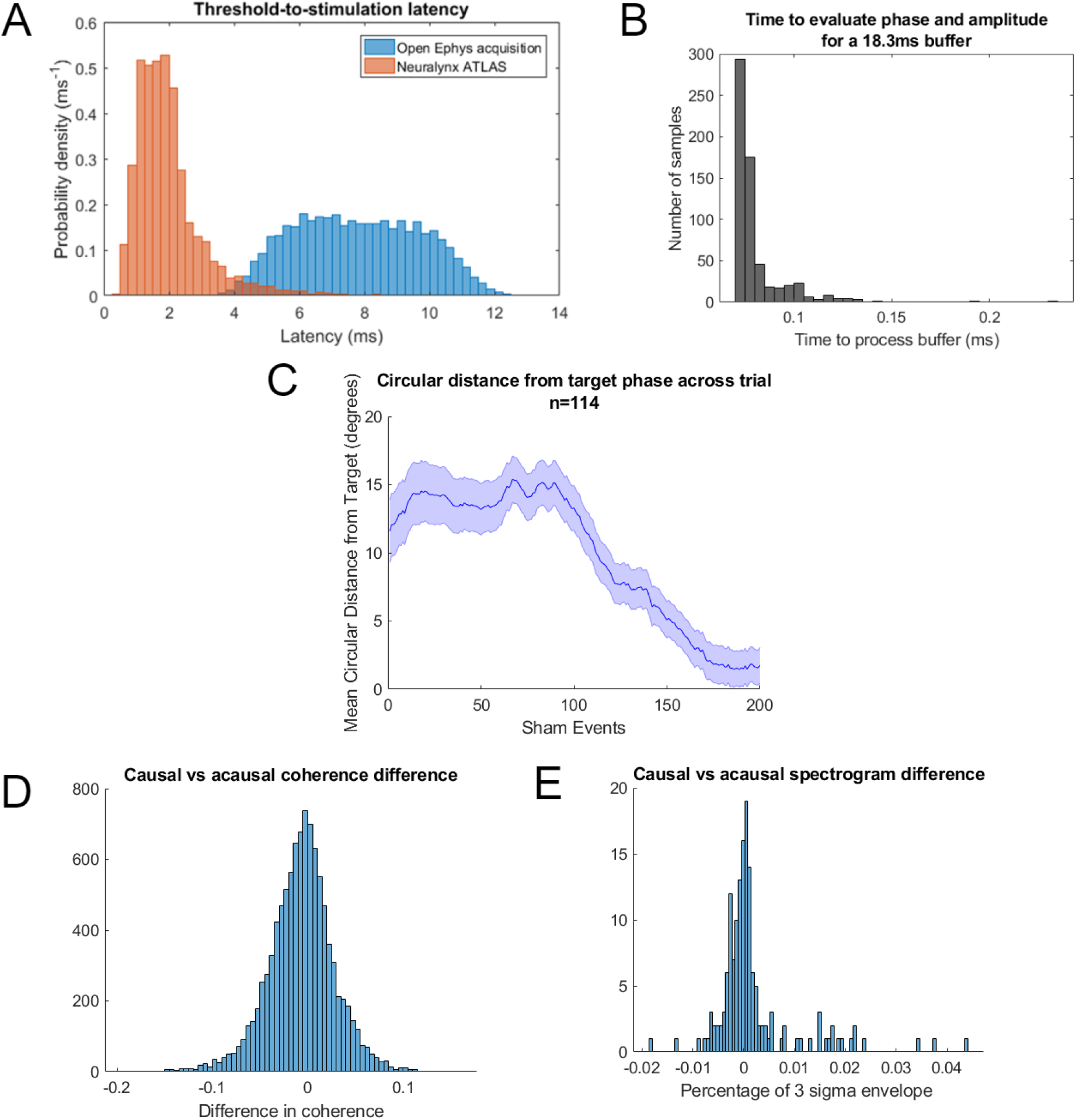
A) The latency between event trigger and perturbation delivery in two widely used acquisition systems. B) The latency between receiving a buffer of data into the algorithm and return of the corresponding analytic signal. C) Phase accuracy improvements over the experiment from the earning algorithm over 114 thirteen minute trials consisting of 200 perturbation pulses. Standard error of the mean is shown as a shaded region. D-E) Histogram displaying the difference between acausa MATLAB and causal C++ implementation of the () coherence and (E) spectrogram of a single channel across 40 frequencies.

### 3.5. Learning Algorithm

TORTE uses a learning algorithm to improve the accuracy of its phase targeting in the presence of system latency, estimation errors and phase bias. Fig. 8C shows phase-locking performance over time in our saline preparation, again targeting 180° in the theta (4-8 Hz) band within the selected IL data channel. Over the first 200 events, accuracy improves by >10°, then stays consistent for the remainder of the experiment. As the learning rate approaches zero during the experiment, the standard deviation decreases.

### 3.6. Real-time Feedback

TORTE’s real-time visualizations closely approximate acausal calculations of the same signals. The mean difference of the coherence output is 0.0065 with a SD of 0.0258 as shown in Fig. 8D. The scale of coherence values are 0 to 1 so these differences can be considered negligible. The difference in the spectrogram causal and acausal measurements were calculated as a percentage of the envelope of the data. The mean difference of the spectrogram is 0.0012% and a SD of 0.0064% as shown in Fig. 8E. These represent small percentages of the full scale, demonstrating the validity of these visualizers for real-time experimental performance tracking.

## 4. Discussion

### 4.1. TORTE

We have presented TORTE, a toolkit to enable scientists to easily implement closed-loop experiments based upon oscillatory activity within continuous neural data. This fills a resource gap by providing an open-source toolkit that can readily be adapted into most existing experimental systems. Further, TORTE is a sub-component of the larger open-source framework of OEGUI. OEGUI takes a modular approach, where plugins can be separately created and compiled without dependence on the main package maintainer. Thus, TORTE leverages other labs’ work to create plugins that stream in data from several commonly used neural acquisition systems.

Although TORTE is a complete toolkit, the plugin architecture also allows TORTE to be extended upon by other plugins that provide additional functionality within OEGUI. An example would be using a behavior to gate the presentation of oscillation informed perturbations. A video tracking plugin could pause the event output of TORTE during periods of non-desired behavior such as grooming, and only allow oscillation-informed perturbations during non-grooming periods. On top of the code being freely available with ample documentation, our lab provides support for users implementing TORTE and the Open Ephys team provides support for using OEGUI. Potential applications include locking an event to the 180° phase of a slow frequency component of a LFP recording to modulate synchrony, sending perturbations in response to high gamma power as a proxy for local spiking rate, or stimulating at periods of high power in the beta band to target tremor-related activity.

### 4.2. Limitations

The toolkit presented is easy to use and flexible, but does have limitations. For processing efficiency, both OEGUI and TORTE are developed in C++, which is extremely computationally efficient but not a commonly-used programming language among life scientists. Where possible, we have made TORTE components easily configurable without a direct code rewrite, but extracting the analytic signal from frequency bands that fall outside of the provided configurations requires knowledge of designing digital filters. TORTE utilizes a Hilbert transformer algorithm which works well for many use cases, however other algorithms may better suit some users’ needs. Only a very rudimentary artifact suppression technique is implemented as described, which is sufficient for intermittent locking to low-frequency oscillations, but may not cover all use cases. Artifacts typically show up as a phase reset, jump to zero phase, or a momentary increase in amplitude. More advanced artifact suppression techniques would need to be assessed on a case by case basis according to recording technique wherein each will include unique artifacts such as blinks, stimulation, etc. Finally, although the toolkit is free and open-source, the user still needs to “assemble” the experiment themselves including the output logic for perturbation presentation. The TORTE team can assist in this process, but may not be able to provide the same level of support that a private company may provide for its products. Several neural acquisition systems are supported, but there are many that OEGUI cannot receive data from yet. As shown, the toolkit is compatible and initial testing has been completed with human invasive and non-invasive systems. It is suitable for basic science experiments, but TORTE is not currently suitable for clinical research, as it has not undergone FDA-compatible design controls.

### 4.3. Algorithm Comparison

As shown in Results, the TORTE Hilbert transformer algorithm provides improved phase accuracy compared to the standard algorithm included in OEGUI. It also provides real time amplitude information which the standard algorithm does not. The Hilbert transformer algorithm works well for many use cases, however other algorithms may better suit some users’ needs. For instance, novel state-space approaches (SSPE) have been proposed as a more principled and reliable way to track oscillations (Wodeyar et al., 2021). TORTE is extendable to new algorithms (we have implemented an initial version of that state-space approach), but each new approach would need to be converted into C++, which is not trivial. The Hilbert transformer algorithm is preferred when the data is narrow band and well characterized or the user wants to implement the simplest solution. The novel SSPE algorithm should improve performance when frequencies of interest are variable or multiple peaks exist within a frequency band. The standard peak/trough detector included in OEGUI may be sufficient in the case of extremely stable, near-sinusoidal oscillations, e.g., occipital eyes-closed alpha.

### 4.4. Future Directions

TORTE is continuously being improved and extended upon, driven by experiments in our own lab as well as our collaborators. The current real-time analytic signal algorithm is fast and reasonably accurate, but oscillatory signal processing is rapidly advancing. For instance, latent variable approaches may estimate and predict oscillatory signals in ways that our current filtering approaches cannot (Yang et al., 2021). We expect to implement these innovations into TORTE as they become available. Similarly, OEGUI itself is rapidly evolving as the Open Ephys platform spreads. A second-generation hardware system will dramatically improve latency by removing USB communication, but will also re-factor OEGUI into the Bonsai architecture. TORTE will be made compatible with these future evolutions, as we intend to adopt them in our own experiments. Future versions may also extend visualization of cross-region oscillatory synchrony to include phase-amplitude coupling or spike-field locking based on real-time spike sorting.

## 5. Conclusion

TORTE provides a platform for rapidly and reproducibly creating oscillation-informed closed-loop experiments. Such experiments are already being implemented, in a preliminary fashion, in areas such as motor rehabilitation, epilepsy, and movement disorders. They are theorized to be applicable to understanding and developing treatments for more complex domains such as mental disorders (Cho et al., 2015).

The availability of a common and flexible toolkit should make these paradigms easier to apply for testing a wide variety of brain functions, accelerating progress in both basic neuroscience and clinical translation.

## Acknowledgments

We would like to thank Dr. Joel Voss, Dr. James Kragel and Sarah Lurie for assistance with generating the Human dataset. We thank Dr. Mo Chen, Dr. Saydra Wilson and Dr. Sarah Olsen for providing the EEG dataset. We thank Dr. Meng-Chen Lo and Rebecca Younk for assistance with generating the Rodent dataset. This work was supported by the Brain & Behavior Research Foundation, Picower Family Foundation, Kent and Liz Dauten Bipolar Disorders Seed Fund at Harvard University, the MnDRIVE Brain Conditions and Medical Discovery Team - Addictions initiatives at the University of Minnesota, and the National Institutes of Health (R21MH109722, R21MH113103, R01EB026938, and R01MH119384).

## 5.1. Author Contributions

M.J.S., E.B.B, and A.S.W. designed the toolkit. M.J.S, E.B.B., and S.S.N. implemented, programmed and preformed analyses on the toolkit. M.J.S. and A.S.W. wrote the paper with input from the other authors. All authors gave final approval to the paper.

